# Cross-Species Dynamics of Transposable Elements in Aging Female Visceral Adipose Tissues

**DOI:** 10.64898/2026.11.13.766407

**Authors:** Syed S Ali, Amrit Gaire, Nissi Paidimukkala, Linshan Laux, Ann V. Hertzel, Mythili Dileepan, Erich Kumerfeld, Paul D. Robbins, Constantin Aliferis, Laura J. Niedernhofer, Jinhua Wang

## Abstract

Transposable elements (TEs) make up nearly half of mammalian genomes and are increasingly implicated in genomic instability, cellular stress responses, aging, and age-related disease. Recent past research has highlighted their significant involvement in disease models such as cancer and Alzheimer’s disease (AD), though much remains to be unraveled regarding these genomic insertions. In this study, we explore the involvement of TEs in aging female mouse and human visceral adipose tissue (VAT) using 10x single-nucleus RNA sequencing (snRNA-seq) data analyzed with the computational tool SoloTE. By assessing the TE-annotated snRNA-seq count matrix using Seurat, we found that retrotransposons—specifically short interspersed nuclear elements (SINEs) and long interspersed nuclear elements (LINEs) on chromosomes 1 and 19—are significantly upregulated in aging female human VAT and in potential senescent cells (*CDKN1A^+^*). Although changes in mouse VAT were less pronounced, long terminal repeats (LTRs) were predominant in aging mouse cells. Furthermore, analysis of upregulated features in aging and *CDKN1A^+^* female human VAT cells revealed that LINE and SINE TEs embedded within specific gene loci—most notably *CUX1*, *SOD2*, *C3*, *NAMPT*, *ALDH1A3*, *IL6*, and the non-coding RNA *LOC105375913*—exhibit a positive regulatory relationship with their host genes. Specifically, higher TE expression may directly upregulate host gene expression (e.g., *C3* TEs driving *C3* upregulation). Consistently ranking at the top of differentially expressed feature lists, these TEs act as potential drivers of aging and cellular senescence in female human VAT and can be plausible targets without risk of disruption of multiple biological systems.

## Introduction

Transposable elements (TEs) make up approximately 45% of the human genome and 37% of the mouse genome^1,2^. They are classified as DNA transposons or retrotransposons. Retrotransposons comprise long terminal repeat (LTR) elements and non-LTR long and short interspersed nuclear elements (LINEs and SINEs). Once dismissed as “junk” DNA, TE-derived sequences are now recognized as a major source of promoters, enhancers and non-coding RNAs that shape host gene regulation^3,4^. The active TE repertoire differs between species. Mice retain active LINE-1 (L1) elements and endogenous retroviruses (ERVs), whereas in human autonomous activity is restricted to L1, which also mobilizes Alu and SVA elements^5^.

In somatic cells, TEs are silenced by heterochromatin and DNA methylation, and these controls erode with age^5,6^. Senescent cells show increased chromatin accessibility at Alu, SVA and L1 elements and accumulate TE transcripts^7^. In late senescence, L1-derived cytoplasmic cDNA triggers a type I interferon response that reinforces the senescence-associated secretory phenotype (SASP). Inhibiting L1 reverse transcriptase attenuates age-associated inflammation in mice^8,9^. Reactivated ERVs similarly reinforce senescence^10^, and TE suppression extends lifespan in *Drosophila*^11^.Whether TE activation causes or accompanies aging remains unresolved.

Visceral adipose tissue (VAT) is a central driver of age-related metabolic decline. Aging VAT shows impaired adipogenesis, immune remodeling and chronic inflammation, which together promote insulin resistance^12,13^. Senescent cells accumulate in VAT, and clearing them restores metabolic function^14,15^. Cells highly expressing *Cdkn1a* (p21) constitute a distinct senescent population whose elimination improves insulin sensitivity^16^. Adipose aging also differs between the sexes, yet females remain underrepresented in mechanistic aging studies^17,18^. Despite the established links between TEs and senescence, TE expression in aging VAT has not been examined.

Single-nucleus RNA sequencing (snRNA-seq) captures mature adipocytes and has enabled cellular atlases of human and mouse adipose tissue^19^. Standard pipelines, however, discard multimapping reads and therefore overlook most TE-derived transcripts^20^. SoloTE overcomes this limitation by quantifying TE expression at individual-locus resolution in single-cell data^21^.

Here, we reanalyzed published snRNA-seq data from young and old female mouse and obese human VAT. Using these data, we characterized age-associated TE expression at the family, locus and cell-type levels, identified differentially expressed TEs within senescence-associated genes and examined TE expression in *CDKN1A* senescent-like cells. These analyses define the TE landscape of aging VAT and identify candidate TE biomarkers of adipose aging and senescence.

## Results

### Assessment of Differentially Expressed Aging TEs at Class, Family & Chromosome Level

The feature matrix annotated with TEs features retrieved via SoloTE for female mouse visceral adipose tissue (VAT) comprised 2,124,480 TEs and 32,206 genes. Differential expression features (DEF) analysis using Seurat identified 414 differentially expressed TEs: 329 down-regulated and 85 up-regulated (Figure 1A). Chromosomal localization of up-regulated TEs revealed marked enrichment of LINE elements on chromosome X, as well as LTR elements on chromosomes 11 and 17 (Figure 1A). Furthermore, LTR-class elements (specifically the ERVK family (Figure 1B) constituted the majority of the 85 up-regulated TEs, whereas L1-LINE elements were predominantly down-regulated on chromosome 8 (Figure 1A).

**Figure 1.**
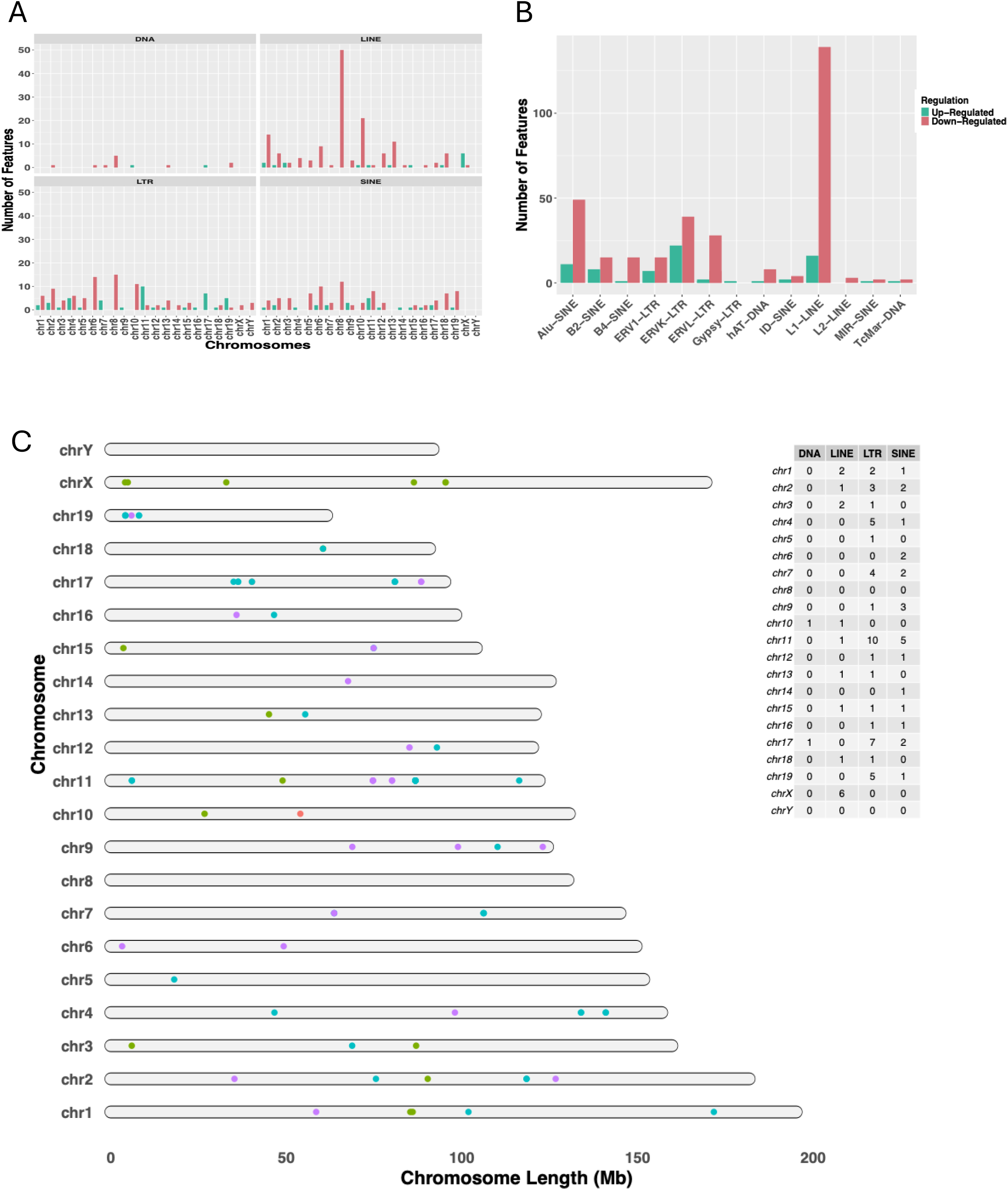
Composition and chromosomal distribution of significantly differentially expressed transposable elements (TEs; 0.05) in female mouse visceral adipose tissue (VAT) during aging. **(A)** Composition of up- and down-regulated TEs identified between old and young cells, categorized by class and chromosome. **(B)** Composition of differentially expressed TEs identified between old and young cells, categorized by TE family. **(C)** Schematic chromosome plot illustrating the genomic locations of up-regulated TEs across mouse chromosomes 1–19, X, and Y. The accompanying table on the right summarizes the total count of up-regulated TEs by class on each chromosome.

Analysis of TE expression in VAT from female humans with obesity demonstrated a more pronounced overall effect. A total of 3,529,980 TEs and 40,943 genes were quantified, with DEF analysis revealing 1,154 down-regulated and 610 up-regulated TEs (Figure 2A). Using the complete nomenclature provided by SoloTE, we mapped chromosomal distribution patterns to determine regional impacts. Alu-SINE and L1-LINE elements showed the highest degrees of up-regulation (Figure 2B). At the chromosomal level, SINE elements exhibited the strongest up-regulation on chromosomes 1, 14, 19, and 20, whereas LINE elements were most up-regulated on chromosomes 14 and 19 (Figure 2A). Conversely, SINE down-regulation primarily affected chromosomes 5, 6, 9, and 12, while LINE down-regulation was localized to chromosomes 3, 6, and 9 (Figure 2A). DNA and LTR elements showed only slight enrichment of up-regulation relative to down-regulation on chromosomes 1 and 19 (Figure 2A).

**Figure 2.**
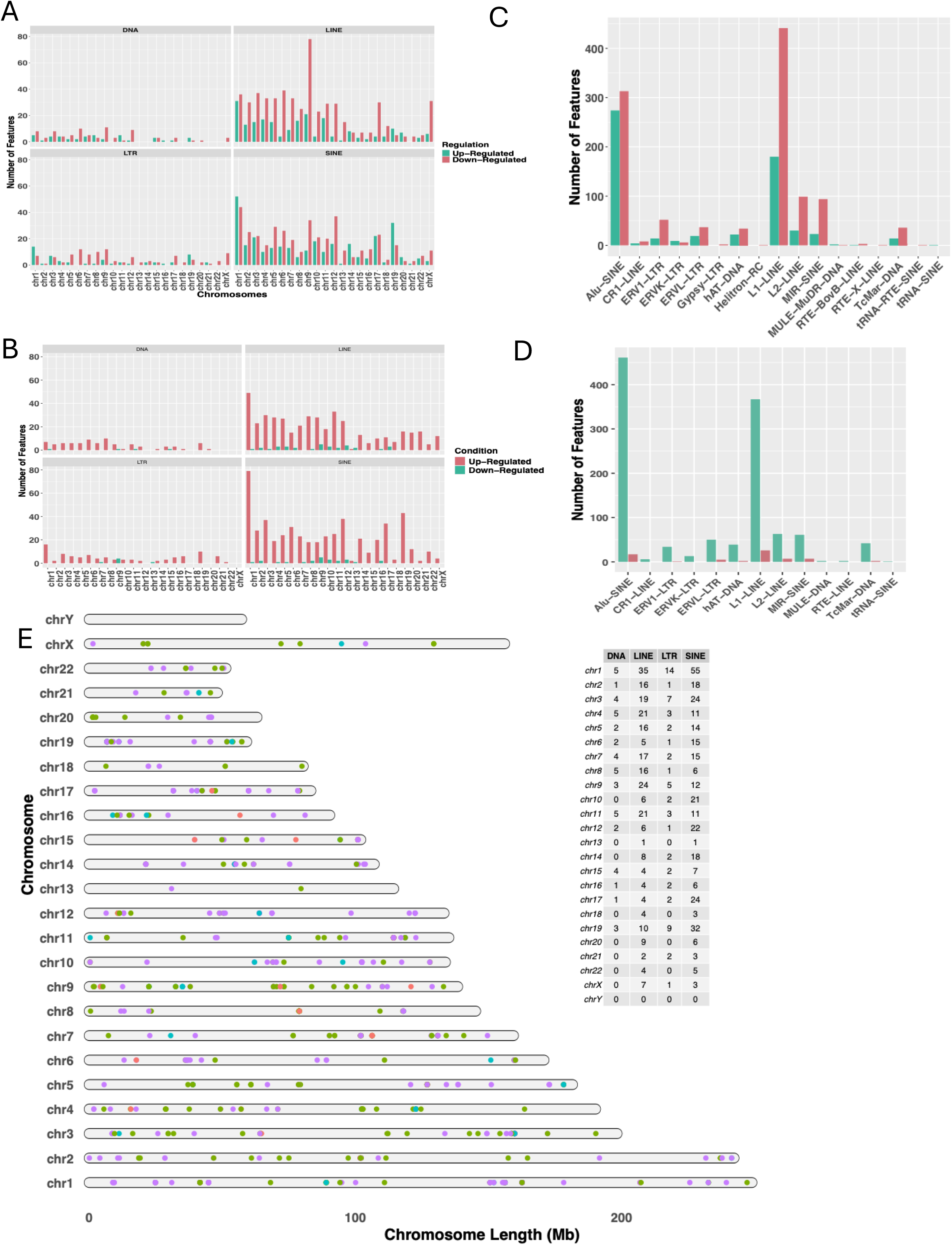
Composition and chromosomal distribution of significantly differentially expressed transposable elements (TEs; 0.05) in female human visceral adipose tissue (VAT) across aging and senescence. **(A)** Composition of up- and down-regulated TEs identified between old and young cells, categorized by class and chromosome. **(B)** Composition of differentially expressed TEs identified between old and young cells, categorized by TE family. **(C)** Composition of up- and down-regulated TEs identified between *CDKN1A*-positive and *CDKN1A*-negative cells, categorized by class and chromosome. **(D)** Composition of up- and down-regulated TEs identified between *CDKN1A*-positive and *CDKN1A*-negative cells, categorized by TE family. **(E)** Schematic chromosome plot illustrating the genomic locations of age-associated up-regulated TEs (old vs. young cells) across human chromosomes 1–22, X, and Y. The accompanying table on the right summarizes the total count of up-regulated TEs by class on each chromosome.

Overall, pseudo bulk analysis indicated that SINE and LINE elements remain more profoundly activated in human aging VAT relative to other TEs, whereas mouse VAT displayed a subtle increase in LTR activity. Finally, comparative expression analysis of *CDKN1A* senescent-like cells with non-senescent cells revealed minimal TE expression in female mouse VAT, whereas female human VAT showed consistent TE expression across all chromosomes, with Alu-SINE and L1 elements exhibiting the greatest up-regulation on chromosomes 1 and 19 (Figure 2C, 2D).

### Characterization of Cell Clusters in female mice and human VAT

Single-nucleus RNA-seq (snRNA-seq) counts annotated with transposable elements (TE) and gene features obtained via SoloTE were analyzed in Seurat to construct cell clusters. Clusters were annotated using reference markers from the human and mouse adipose tissue atlas published by Emont et al. (2022). In the mouse VAT, we identified ten cell types: adipocytes, adipose stem and progenitor cells (ASPCs), B cells, endothelial cells, macrophages, mesothelial cells, monocytes, pericytes, smooth muscle cells (SMCs), and T cells (Figure 3A). Among these, ASPCs, B cells, endothelial cells, macrophages, monocytes, pericytes, SMCs, and T cells showed slight increases in abundance in aged tissues, though the differences were modest (Figure 3B).

**Figure 3.**
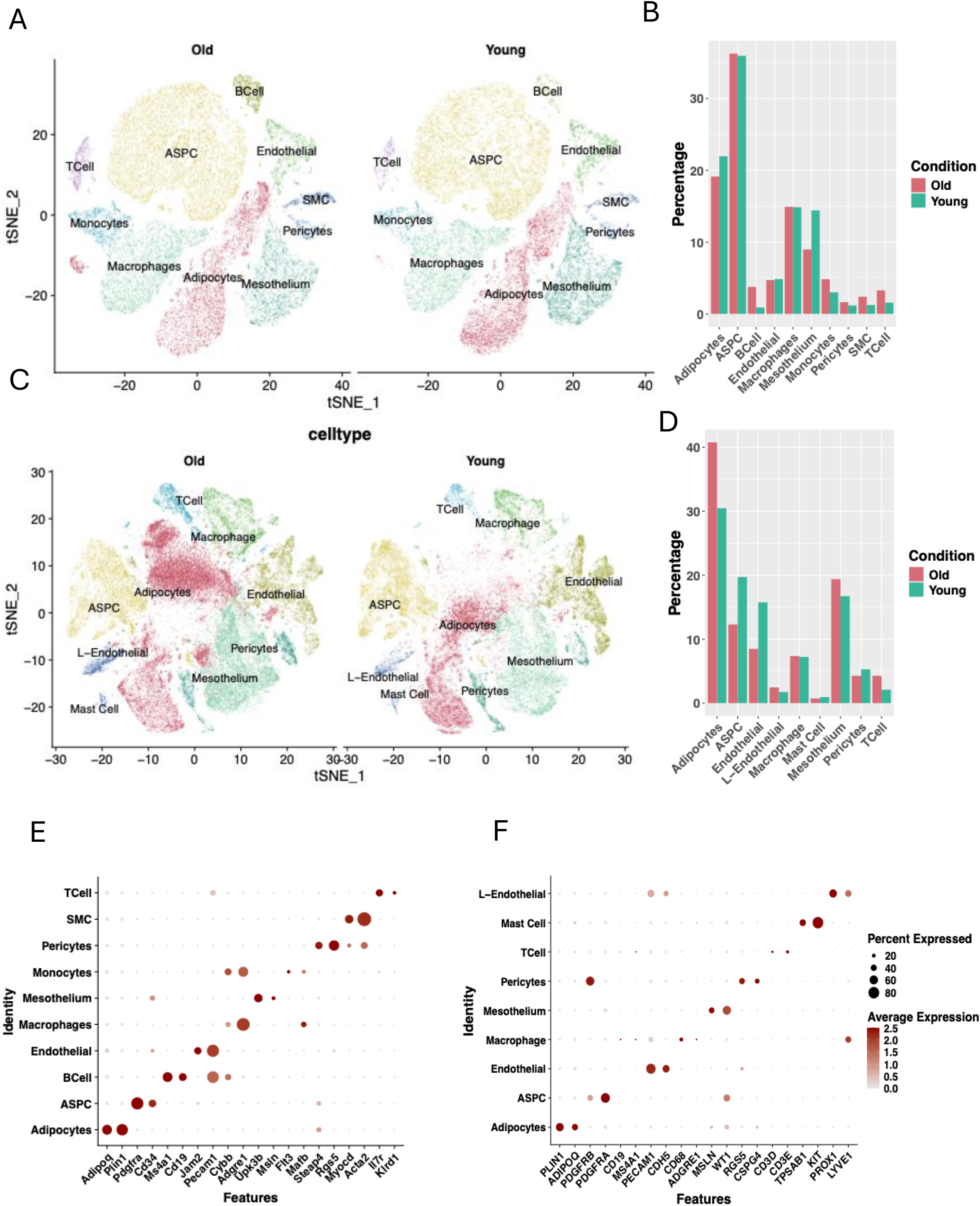
Single-cell characterization and age-dependent cellular composition of female mouse and human visceral adipose tissue (VAT). **(A)** t-SNE plot of female mouse VAT illustrating identified cell populations: adipocytes, adipose stem and progenitor cells (ASPCs), B cells, endothelial cells, macrophages, mesothelial cells, monocytes, pericytes, smooth muscle cells (SMCs), and T cells. **(B)** Proportions of each cell type across age conditions in female mouse VAT. **(C)** t-SNE plot of female human VAT *(note: verify species—your draft says “mouse” here, though panels C–D pair with human VAT)* illustrating identified cell populations: adipocytes, ASPCs, endothelial cells, lymphatic endothelial cells (L-Endothelial), macrophages, mast cells, mesothelial cells, pericytes, and T cells. **(D)** Proportions of each cell type across age conditions in female human VAT. **(E)** Dot plot showing canonical marker genes used to annotate cell clusters in female mouse VAT. **(F)** Dot plot showing canonical marker genes used to annotate cell clusters in female human VAT.

For female human obesity VAT, we identified adipocytes, ASPCs, lymphatic endothelial cells (L-Endothelial), macrophages, mast cells, mesothelial cells, pericytes, and T cells (Figure 3C). Cell proportion analysis between young and old tissue conditions revealed elevated numbers of adipocytes, mesothelial cells, macrophages, and T cells in aged samples (Figure 3D). The marker genes used to define these cell type annotations are displayed in the Seurat dot plots (Figures 3E and 3F).

### Assessment of Up-regulated TEs at the Gene and Cellular Level in female mice VAT

Average expression of up-regulated transposable elements (TEs) annotated to known genes— identified via pseudo bulk differential analysis comparing old and young tissue—is presented in the heatmap in Figure 4A. The analysis revealed that most aging-associated TEs were predominantly expressed in immune cell populations, specifically B cells, macrophages, and T cells. Among non-immune lineages, the endothelial cell cluster demonstrated the highest levels of TE expression.

**Figure 4.**
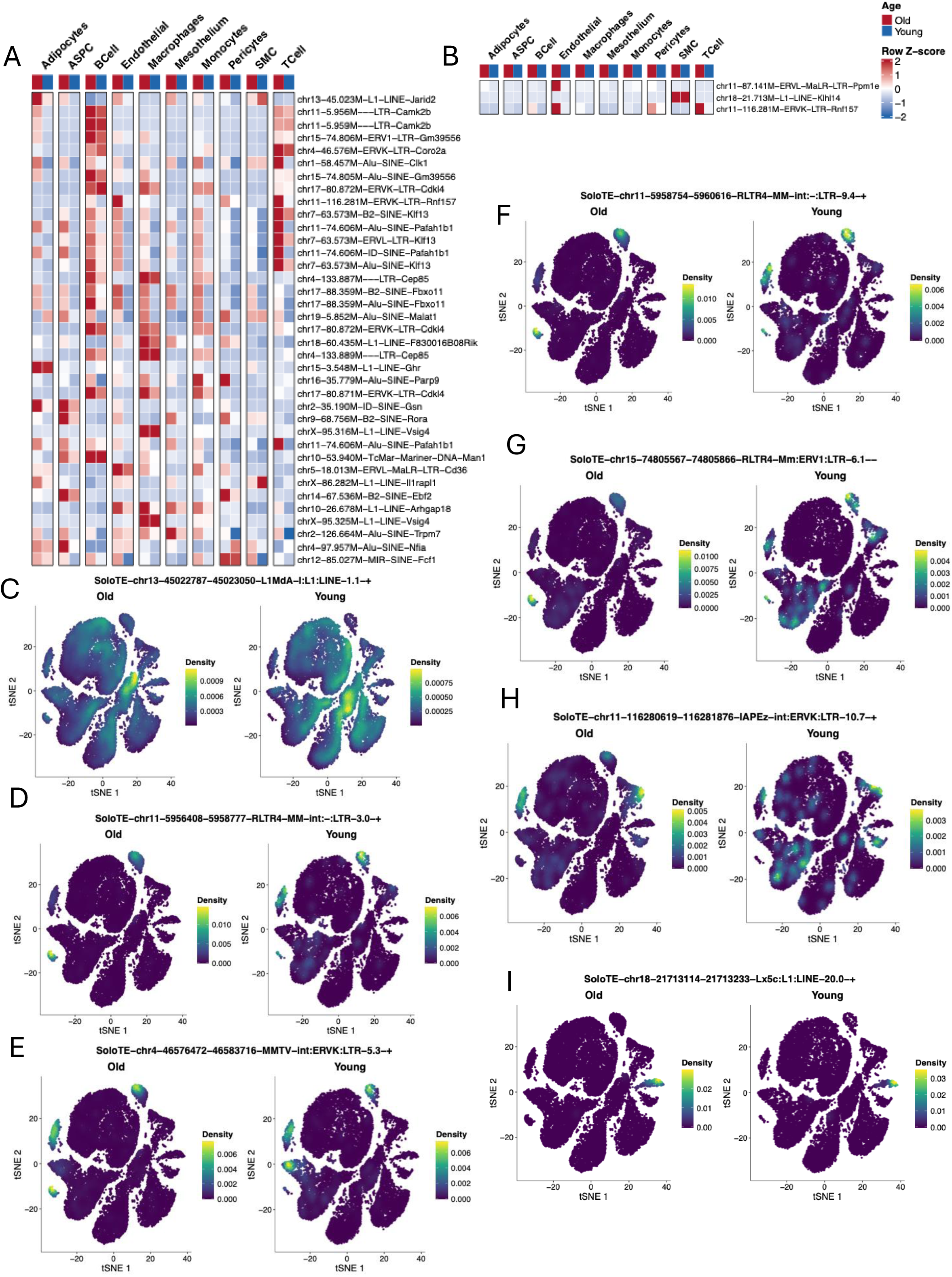
Cell-type-specific expression of gene-overlapping transposable elements (TEs) across age and *Cdkn1a* status in female mouse visceral adipose tissue (VAT). **(A)** Heatmap showing scaled expression levels (−2 to +2) of gene-overlapping TEs identified between old and young cells across cell types and age conditions. Rows indicate individual TEs annotated by chromosome, starting genomic position, family, class, and host gene; columns represent cell types grouped by condition. **(B)** Heatmap showing scaled expression levels (−2 to +2) of gene-overlapping TEs identified between *Cdkn1a*-positive and *Cdkn1a*-negative cells across cell types and conditions, annotated as in (A). **(C–I)** Density plots illustrating the expression distributions of selected TEs in old (left) versus young (right) cells, with the full TE identifier labeled above each plot.

These up-regulated TEs were primarily located within sequences corresponding to *Kif13*, *Cdk4*, *Pafah1b1*, *Camk2b*, *Cap85*, *Pbx11*, and *Vsig4*. Further examination using density plots (Figures 4C–4I) for seven of the top TEs mapped to *Camk2b* and *Jarid2* highlighted adipocyte and B-cell clusters as primary sites of interest. Finally, comparative analysis between *Cdkn1a* senescent-like cells and non-senescent cells identified only three annotated TEs—located within *Ppm1e*, *Klhl14*, and *Rnf157*—which were predominantly expressed in aged endothelial cells (Figure 4B). *Kif13a* and *Pafah1b1* were downregulated, whereas *Cdk4*, *Camk2b*, *Vsig4*, and *Jarid2* were upregulated. *Cap85* and *Pbx1* did not meet the predefined significance threshold (p < 0.05). Additional genes, including *Ppm1e* and *Rnf157*, were also not significantly differentially expressed, while *Klhl14* was upregulated. Overall, the identified genes were associated with diverse biological processes, including intracellular transport, metabolic regulation, inflammatory signaling, and macrophage function. However, their expression patterns did not consistently correspond with previously reported aging- and senescence-associated changes.

### Assessment of Up-regulated TEs at the Gene and Cellular Level in female obese human VAT

Because a greater number of gene-annotated TEs were retrieved from human VAT compared to mouse VAT, these features were evaluated in two distinct stages. First, we examined the top 30 TEs within each major class (SINE, LINE, LTR, and DNA), identifying 54 TEs out of 120 that overlapped with annotated gene sequences (Figure 5A). Assessment of these 54 TEs revealed that the majority were localized within non-coding RNA *LOC105375913*, as well as *HIVEP3*, *ALDH1A3*, *IQCJ-SCHIP1*, *C3*, *NAMPT*, *BBS12*, and *MARCHF3*, with chromosomes 1, 7, 8, 15, and 19 occurring most frequently. DNA and LINE elements overlapped heavily with *LOC105375913*, *HIVEP3, NAMPT*, whereas SINE elements showed strong overlap with *C3*. Cell-type expression profiling demonstrated that these TEs were predominantly expressed in aged adipocytes, ASPCs, and mesothelial cells (Figure 5A).

**Figure 5.**
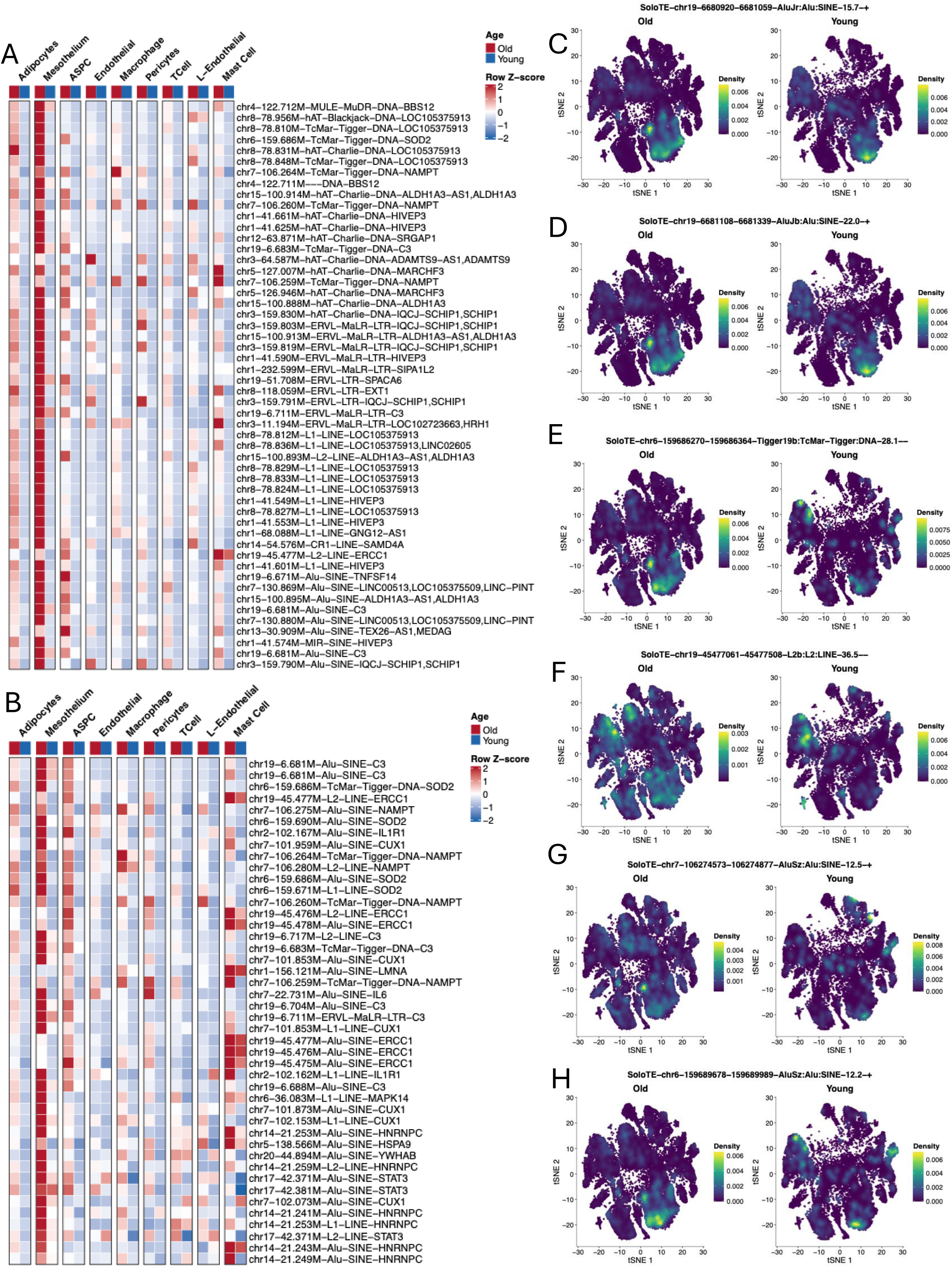
Cell-type-specific expression of age-upregulated, gene-overlapping transposable elements (TEs) in female human visceral adipose tissue (VAT). **(A)** Heatmap showing the expression of 54 gene-overlapping TEs identified from the top 30 age-upregulated TEs (old vs. young cells) per TE class. Rows indicate individual TEs annotated by chromosome, starting genomic position, family, class, and host gene; columns represent cell types grouped by age condition. **(B)** Heatmap showing the expression of all age-upregulated TEs overlapping with senescence-associated genes from the SenMayo and CellAge gene panels. **(C–H)** Density plots illustrating the expression distributions of selected TEs in old (left) versus young (right) cells, with the TE identifier labeled above each plot.

Next, we assessed TEs located within gene sequences associated with cellular senescence, retrieved from the SenMayo and CellAge gene sets (Figure 5B). These TEs showed frequent overlaps with *NAMPT*, *ERCC1*, *C3*, *HNRNPC*, *CUX1*, *SOD2*, and *STAT3*, with chromosomes 6, 7, 14, 17, and 19 being the most represented. SINE elements on chromosome 19 overlapped with *C3*, consistent with the top TE patterns observed in Figure 5A. Similarly, DNA elements on chromosome 7 overlapped heavily with *NAMPT*, mirroring results from the initial 54 TEs (Figure 5A). Additionally, SINE and LINE TEs on chromosome 7 overlapped with *CUX1*, LINE and SINE TEs on chromosome 6 overlapped with *SOD2*, SINE and LINE TEs on chromosome 14 overlapped with *HNRNPC*, and SINE and LINE TEs on chromosome 19 overlapped with *ERCC1* (Figure 5B) – these genes were unique not consistent with results shown in figure 5A. Density plots of Top 6 TEs (Figures 5C–5H) highlighted mesothelial cells as the primary cell cluster of focus, with partial involvement of ASPCs.

To benchmark whether specific TEs possess cell type–specific expression patterns, we performed differential expression analysis across cell types to identify the top TE marker features for each lineage (Figures 6A and 6B). The identification of cell type–specific TE markers established a baseline proof-of-concept that distinct TE features can serve as lineage-specific markers, establishing the rationale to test whether specific TEs similarly mark aging and senescent cell populations. Building on this benchmark, we compared *CDKN1A* senescent-like cells and non-senescent cells and identified the top 40 gene-overlapping TEs (Figure 7A). Chromosomes 6, 7, 8, and 19 accounted for the majority of these top TEs, with sequence overlaps localized primarily to *C3*, *NAMPT*, *SOD2*, and *LOC105375913* (Figure 7A). SINE and LINE elements constituted most of these features, with SINE elements showing the highest representation (Figure 7B). Finally, density plots (Figures 7C–7E) confirmed that mesothelial cells displayed the highest expression of aging based and senescence based TEs among all cell types.

**Figure 6.**
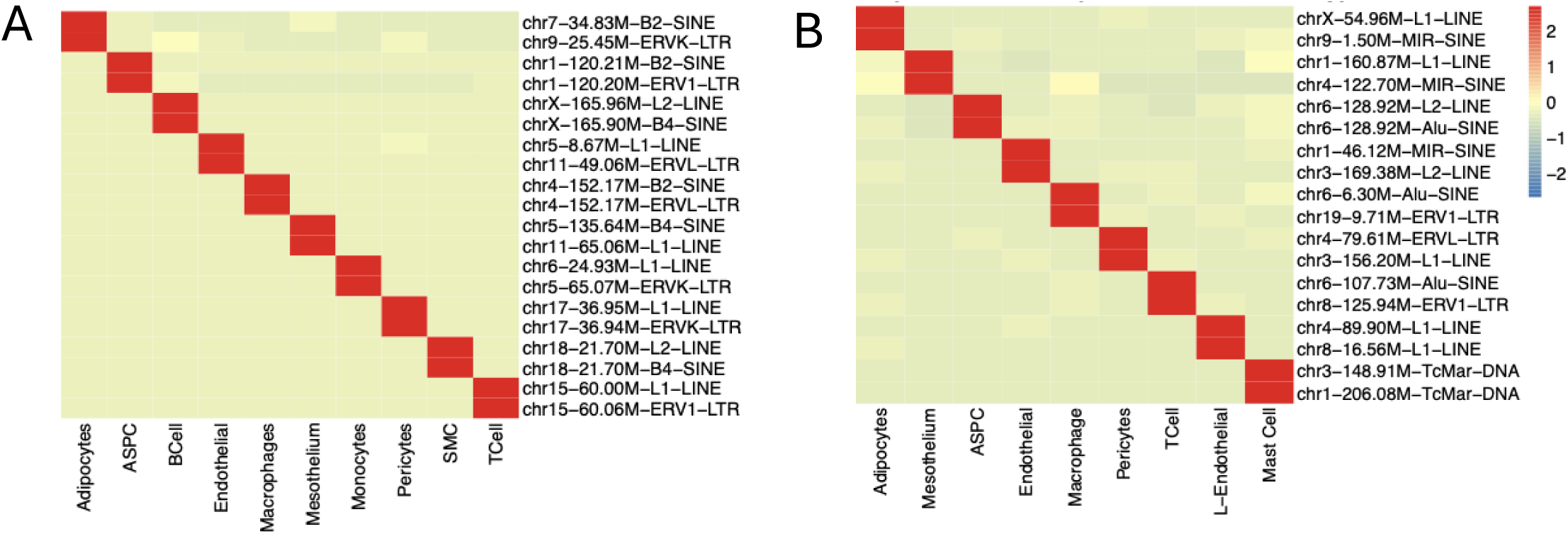
Expression of cell-type-specific transposable elements (TEs) in female mouse and human visceral adipose tissue (VAT). **(A)** Heatmap showing the expression of marker TEs enriched across distinct cell types in female human VAT. **(B)** Heatmap showing the expression of marker TEs enriched across distinct cell types in female human VAT.

**Figure 7.**
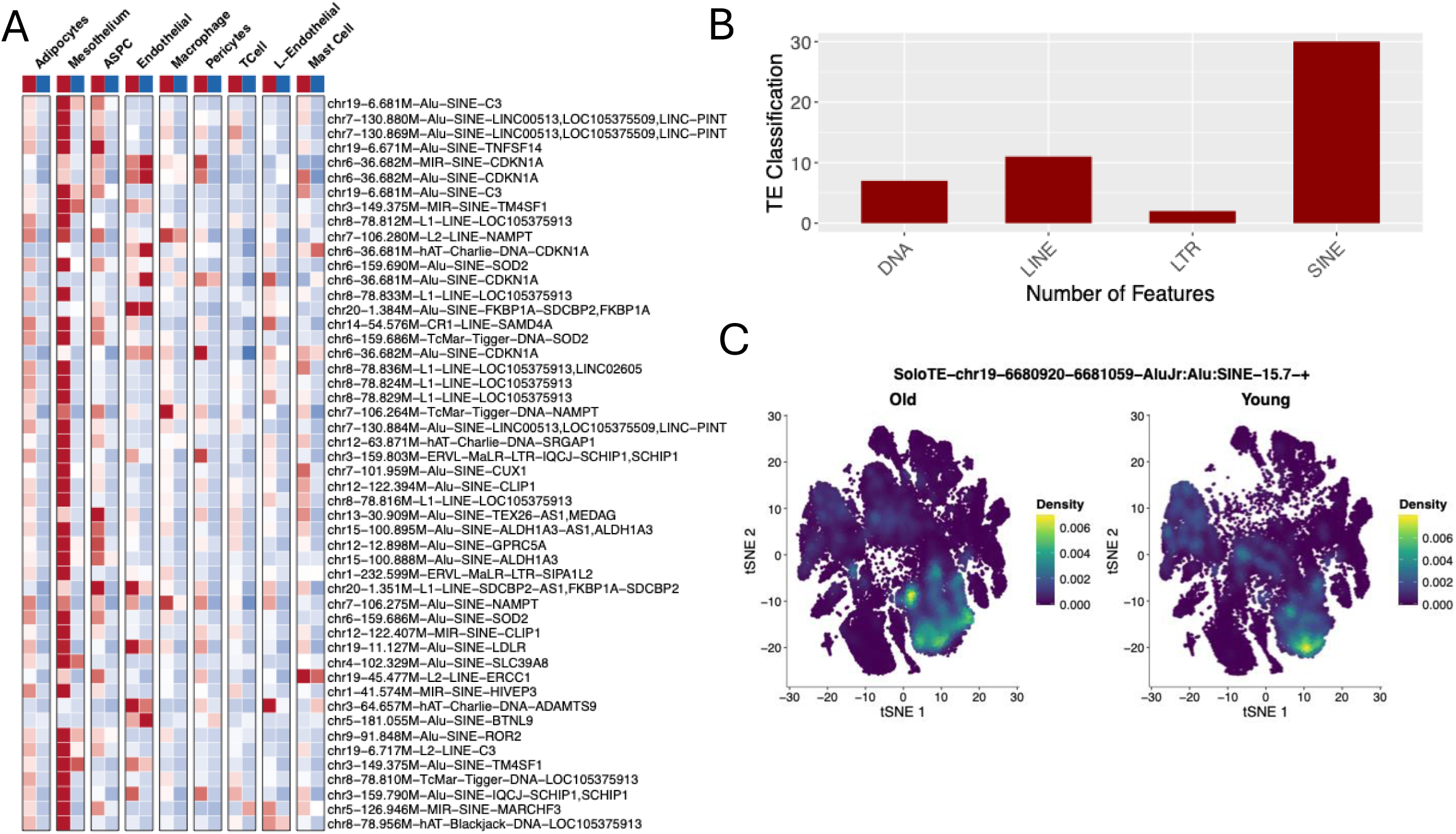
Cell-type-specific expression and class composition of the top 50 *CDKN1A*-associated, gene-overlapping transposable elements (TEs) in female human visceral adipose tissue (VAT). **(A)** Heatmap showing the expression of the top 50 up-regulated, gene-overlapping marker TEs (identified between *CDKN1A*-positive and *CDKN1A*-negative cells) enriched across distinct cell types in female human VAT. **(B)** Class composition of the top 50 marker TEs. **(C** Density plot illustrating the expression distribution of the top marker TE in young (left) versus old (right) cells, where highlighted regions indicate higher expression levels.

## Discussion

In 1990, W. Murray proposed a theory suggesting that transposons increase in activity as organisms age^22^. Subsequent studies have indicated that this phenomenon may also occur in senescent cells. Here, we investigated this hypothesis using female mouse and human visceral adipose tissue (VAT).

From our analysis, we found that LTRs are predominantly expressed in mouse VAT. However, the upregulation of transposable elements (TEs) was not very significant compared to what we observe in humans, and we did not find substantial upregulation of SINE and LINE elements. Nevertheless, among the 85 TEs we identified as upregulated in an age-dependent manner, many were LTRs, confirming that LTRs are expressed. Although we quantified a very large number of TEs (>1,000,000), only a very small fraction is upregulated, which aligns with current literature stating that most remain inactive over time due to evolutionary changes^23^. In addition, finding ERVK-LTRs to be upregulated within our significant TEs derived from DEFs is also a plausible observation. This occurs because as an organism ages, the methylated chromatin structure opens, making TEs active^23^. This mechanism applies to our human VAT analysis as well, which is why we observe TEs upregulated in human VAT. Furthermore, the reason we observe more upregulated TEs in humans versus mice is because we suspect the methylation is decreased more in humans than in mice.

Chromosomes 11, 17, and X, which we found to contain the most upregulated TEs in mice, are linked to aging. A quantitative trait locus on chromosome 11 has been linked to lifespan in mice, with *Pttg1* (Securin) identified as the underlying gene regulating the sensitivity of hematopoietic stem and progenitor cells to replication stress^24^. Chromosome 17 is known to house miR-125a, which also plays a role in aging^25^, while chromosome X has been linked to aging in the brain hippocampus^25^. In humans, although we found multiple involved chromosomes, chromosomes 1 and 19 showed the most upregulation. Chromosome 1 houses the *LMNA* gene, which is related to senescence, though there are no specific studies on the chromosome itself. Two published studies on chromosome 19 claim that chromosome 19 is related to late-onset Alzheimer’s disease^26^, and six loci were found across chromosomes 19, 20, and 13 that influence the age of natural menopause^27^.

Alu-SINE and L1-LINE elements were the two transposable element subfamilies upregulated most in our aging analysis. Substantial evidence links both Alu-SINE and L1-LINE elements to the aging process. Mechanistically, host cell machinery translates L1 RNA into ORF1 and ORF2 proteins^28^. Alu-SINE elements hijack ORF2 to form their own RNPs, enabling them to integrate into the genome and drive genomic instability as well as interferon response activation^28^.

Among the up-regulated 85 TEs in female mouse VAT, 45 that overlapped with known gene sequences, we did not find conclusive evidence regarding their host genes link to tissue aging or cellular senescence. However, this does not imply that the tissue did not undergo age-related changes; rather, our findings suggest that the TEs identified in female mouse VAT were not demonstrably associated with the regulation of aging or senescence-related processes.

We identified 610 transposable elements (TEs) significantly upregulated in aging female human VAT. Among the top 30 TEs per class we retrieved, 54 overlapped with known gene sequences, and 44 TEs from the entire DEFs list overlapped with senescence-associated genes found in SenMayo and CellAge gene panels. Furthermore, analysis of the top 50 differentially expressed TEs found by comparing potentially senescent and non-senescent cells predominantly revealed to be composed of SINE (specifically Alu) and LINE (specifically L1) elements - consistent with existing literature. Cross-referencing the top TEs from both aging and senescent-cell analyses highlighted several host genes with multiple TE overlaps — including *LOC105375913*, *HIVEP3*, *ALDH1A3*, *IQCJ-SCHIP1*, *C3*, *NAMPT*, *BBS12*, *MARCHF3*, *ERCC1*, *HNRNPC*, *CUX1*, *SOD2*, and *STAT3* (Figures 7 and 8). All these host genes were upregulated, indicating a positive relationship between them and their TEs.

Cross-comparison prioritized TEs overlapping *C3* and *NAMPT gene sequences* as prime candidate targets. C3 serves as the central component of the complement cascade and undergoes proteolytic cleavage into C3a and C3b^28^. Upregulation of C3a and its receptor (C3aR) drives C3a–C3aR signaling, which promotes cellular senescence in mouse fibroblasts^29^. However, direct inhibition of C3 may be counterproductive, as previous studies suggest C3 depletion can attenuate senescent cell sensitivity to senolytic agents^30^. Consequently, targeting TE-mediated regulation of *C3* may offer a mechanism to modulate senescence pathways without globally compromising systemic complement function. Similarly, *NAMPT* represents a compelling candidate; as the rate-limiting enzyme in NAD+ biosynthesis, NAMPT converts nicotinamide to nicotinamide mononucleotide^31^. While intracellular NAD+ levels decline during senescence despite elevated intracellular NAMPT expression, senescent cells actively secrete extracellular NAMPT (eNAMPT), propagating pro-inflammatory responses (12). Given this link, targeted modulation of *NAMPT*-associated TEs may mitigate senescent phenotypes while avoiding the systemic off-target effects typical of complete gene knockout.

### Conclusion

In conclusion, our findings highlight a fundamental interspecies difference: aging and senescent mouse VAT lacks significant TE activity compared to human VAT. In humans, TE expression during VAT aging is dominated by Alu-SINE and L1-LINE retrotransposons. Rather than age associated TEs concentrating at specific loci such as chromosomes 1 and 19, we find up-regulation may occur broadly across multiple chromosomes. Cross-referencing age groups with marker-defined senescent states (□□□□1□^+/-^) specifically pinpoints TEs overlapping the *C3* and NAMPT gene locus. These *C3* and NAMPT associated TEs represent promising diagnostic markers and candidate targets for therapeutic anti-senescence strategies.

## Methods

### Collection of Tissue Data

snRNA-seq samples from female mice aged 6 months and 24 months and female humans diagnosed with obesity were obtained from age groups 18-35 (young) and 52-75 (old) from the Institute of Biology of Aging and Metabolism at the University of Minnesota. The snRNA-seq samples were generated using the 10x Genomics 5’ chemistry and 3-6 samples were included in each age group in mice and human VAT.

### Processing and Quantification of Single-Nucleus RNA-seq Data and Locus-Level Transposable Elements

Raw FASTQ files from 10x Genomics single-nucleus RNA sequencing (snRNA-seq) were processed using Cell Ranger (v9.0)^32^. Sequence reads were aligned to the pre-built reference genomes provided by 10x Genomics: GRCm39 for mouse samples and GRCh38 for human samples. The resulting aligned BAM files were subsequently used as input for SoloTE^21^ to generate combined gene and transposable element (TE) count matrices. To retain high-quality cells, the SoloTE matrices were filtered based on valid cell barcodes identified by Cell Ranger (located in the filtered feature-barcode matrix output directory for each sample). Matrix filtering and subsetting were performed using Seurat (v5.0)^33^ in R (v4.4.0)^34^. Specifically, locus-level TE counts were derived from the SoloTE loci/ output directory, which contains feature, barcode, and count matrices corresponding to transposable elements resolved at individual genomic loci.

### Quality Control, Integration, and Clustering of Transposable Element-Annotated Expression Data

Following preliminary cell-barcode filtering using Cell Ranger, raw count matrices were subjected to standard downstream quality control (QC) and analysis within the Seurat workflow. Cells were filtered based on the number of detected genes (nFeature_RNA) and mitochondrial gene content; specifically, cells were retained if they contained between 200 and 5,000 detected genes and less than 5% mitochondrial read counts per sample. Expression counts for each sample were log-normalized using the NormalizeData() function in Seurat. Subsequently, the top 2,000 highly variable features were identified per sample using FindVariableFeatures(). Data integration across samples was performed using a representative reference sample. The integrated assay was scaled, and principal component analysis (PCA) was performed. Dimensionality reduction via Uniform Manifold Approximation and Projection (UMAP) and nearest-neighbor graph construction were executed based on the first 15 principal components. Graph-based clustering was conducted at a resolution of 0.5, and cell clusters were manually annotated using marker genes established by Emont et al. ^35^.

### Identification, Genomic Annotation, and Senescence Association of Age-Dependent Transposable Elements

To identify age-dependent transposable elements (TEs), a pseudobulk differential expression analysis was performed comparing old and young cells in female mouse and human visceral adipose tissue (VAT) datasets using the FindMarkers() function in Seurat. TEs were identified as features prefixed with “SoloTE” that exhibited a false discovery rate (FDR)-adjusted *p*-value < 0.05. The embedded feature nomenclature—containing chromosome location, genomic start/end coordinates, sub-family, family, and class—enabled visualization at the chromosome, class, and family levels. Overlapping gene sequences were mapped to genomic coordinates using the GenomicRanges^36^ R package. To identify senescence-associated TEs, TEs overlapping with established senescence gene sets (SenMayo^37^ and CellAge^38^) were identified. A similar workflow was applied to compare *Cdkn1a^+^* (*p*21^+^) and *Cdkn1a^-^* cells across both species to confirm age-associated TEs and furthermore senescence associated TEs.

### Visualization of Age-Associated Transposable Element Expression

Directionality of TE expression (upregulation or downregulation during aging) was determined based on log2 fold-change values. Data visualizations were generated using the ggplot2 R package^39^, and heatmaps were constructed using the pheatmap R package^40^. Density plots were created using Nebulosa package (V3.23)^41^.

## Funding Acknowledgments

The work reported here received financial support from NIH grants U54 AG079754 and U54 AG076041. The UMN Masonic Cancer Center is supported by the U.S. National Institute of Health and National Cancer Institute [Cancer Center Support Grant CA-77598].

## Author Contributions

J.W., L.J.N., and P.D.R. conceived and designed the study. S.A., A.G., and N.P. carried out the data collection and the data analysis. N.P., L.L., A.H., M.D., E.K., C.A., P.D.R., L.J.N., J.W. contributed to data interpretation. S.A., A.G., and J.W. wrote the manuscript, and J.W. reviewed and edited the manuscript with feedback from all the authors. All authors reviewed and approved the final manuscript.

